# Cyclic Electromagnetic DNA Simulation (CEDS) targeting Telomere Repeat Sequence can enhance Anticancer Effect, while CEDS targeting Canonical E-box Sequence induce Oncogenic Effect in Cells

**DOI:** 10.1101/2024.08.25.609549

**Authors:** Yeon Sook Kim, Suk Keun Lee

## Abstract

Cyclic electromagnetic DNA simulation (CEDS) was used in the previous study (*1*) to stimulate the base pair polarities of oligo-dsDNAs, plasmid DNAs, and miRNAs. This study applied dodecagonal CEDS to target telomere repeat sequence (TTAGGG*-CEDS), canonical E-box sequence (CACGTG*-CEDS) in genomic DNA of RAW 264.7 cells and KB cells. During cell culture, CEDS was performed at 20-25 Gauss for 20 min, and followed by cytological observation and immunoprecipitation high-performance liquid chromatography (IP-HPLC) using 350 antisera. KB cells treated with TTAGGG*-CEDS showed a significant decrease in cell density and metachromasia in toluidine blue staining compared to untreated control, while only a slight decrease in cell density and metachromasia was observed by telomere mutation sequence, TTTGGG*-CEDS. Conversely, CACGTG*-CEDS induced almost no change in cell density, cell number, metachromasia in toluidine blue staining compared to random sequence*-CEDS (2(ACGT)*-CEDS) and untreated control. IP-HPLC results showed that TTAGGG*-CEDS induced a potent anticancer effect by downregulating RAS, oncogenesis, and telomere signaling, and activating NFkB signaling-p53-and FAS-mediated apoptosis-innate and cellular immunity signaling axis compared to untreated control and positive control performed with nonspecific poly-A 12A*-CEDS. In contrast, CACGTG*-CEDS exhibited a significant oncogenic effect by enhancing of RAS and NFkB signaling and resulting oncogenesis-glycolysis-chronic inflammation signaling axis. Therefore, it is suggested that CEDS is able to target DNA motif sequences in genomic DNA, and that TTAGGG*-CEDS and CACGTG*-CEDS are favorable candidates to regulate important genes involved in oncogenesis, inflammation, apoptosis, etc.

## Introduction

The previous study showed that CEDS increased DNA hybridization potential and influenced the functions of six bps double-stranded DNA (dsDNA), 24 bps dsDNAs, plasmid DNAs, and miRNAs through *in vitro* experiments (*1*). In the same line, this study demonstrated CEDS targeting DNA motif sequences in genomic DNA. To increase the efficiency of CEDS, this study employed both strands of DNA motif sequences, which were designated as “target sequence*-CEDS.”

Telomere repeat sequences facilitate the replication of chromosome ends and the generation of a T-loop with a telomere-binding protein complex, shelterin, to prevent chromosomal degradation. Telomere shortening and instability can induce aging and various diseases due to telomere dysfunction and chromosomal instability (*2*). Furthermore, they can form G-quadruplex structure with external Hoogsteen bonding strength (*3*), which plays an important role in chromatin protection, DNA replication, transcription, and genomic stability. In this study, a telomere repeat sequence, dsTTAGGG, was selected for CEDS.

The E-box binding proteins play a pivotal role in regulating transcription activity for genome-wide transcription of numerous genes. Their numbers and types vary depending on the binding specificity of E-box (*4*). Among the consensus sequences of E-box, dsCANNTG, the canonical E-box sequence dsCACGTG, which is known to be preferentially bound by cMyc, was selected in this study for CEDS to know the transcriptional changes of cMyc target genes.

### Cytological observation of KB cells treated with CEDS using a telomere repeat sequence

The KB cells derived from human epithelial cell (ATCC, USA) were cultured on culture dishes (SPL, Korea) in Dulbecco’s modified Eagle’s medium (WelGene Inc., Korea) supplemented with 10% fetal bovine serum (WelGene Inc., Korea), 100 units/mL carbenicillin, 100 μg/mL streptomycin, and 250 ng/mL amphotericin B (WelGene Inc., Korea), in 5% CO_2_ incubator at 37.5°C. Approximately 50% confluent KB cells grown on culture dish surfaces were treated with dodecagonal CEDS using a telomere repeat sequence*-CEDS (TTAGGG*-CEDS) at 23-25 Gauss for 20 min twice a day. Following three days of experiments, the cells were fixed with 10% buffered formalin and stained with hematoxylin and eosin (H&E).

TTAGGG*-CEDS exhibited a notable reduction in cell density on the culture dish as observed visually, and a comparable decline in the number of KB cells as observed microscopically, in comparison to untreated control and CEDS using a random sequence 2(ACGT)*-CEDS. In the high-magnification, the cells treated with TTAGGG*-CEDS exhibited notable reduction in cell number, accompanied with frequent pyknotic cells undergoing apoptosis compared to the cells not treated with TTAGGG*-CEDS and the cells treated with 2(ACGT)*-CEDS (Fig. 1).

**Figure 1.**
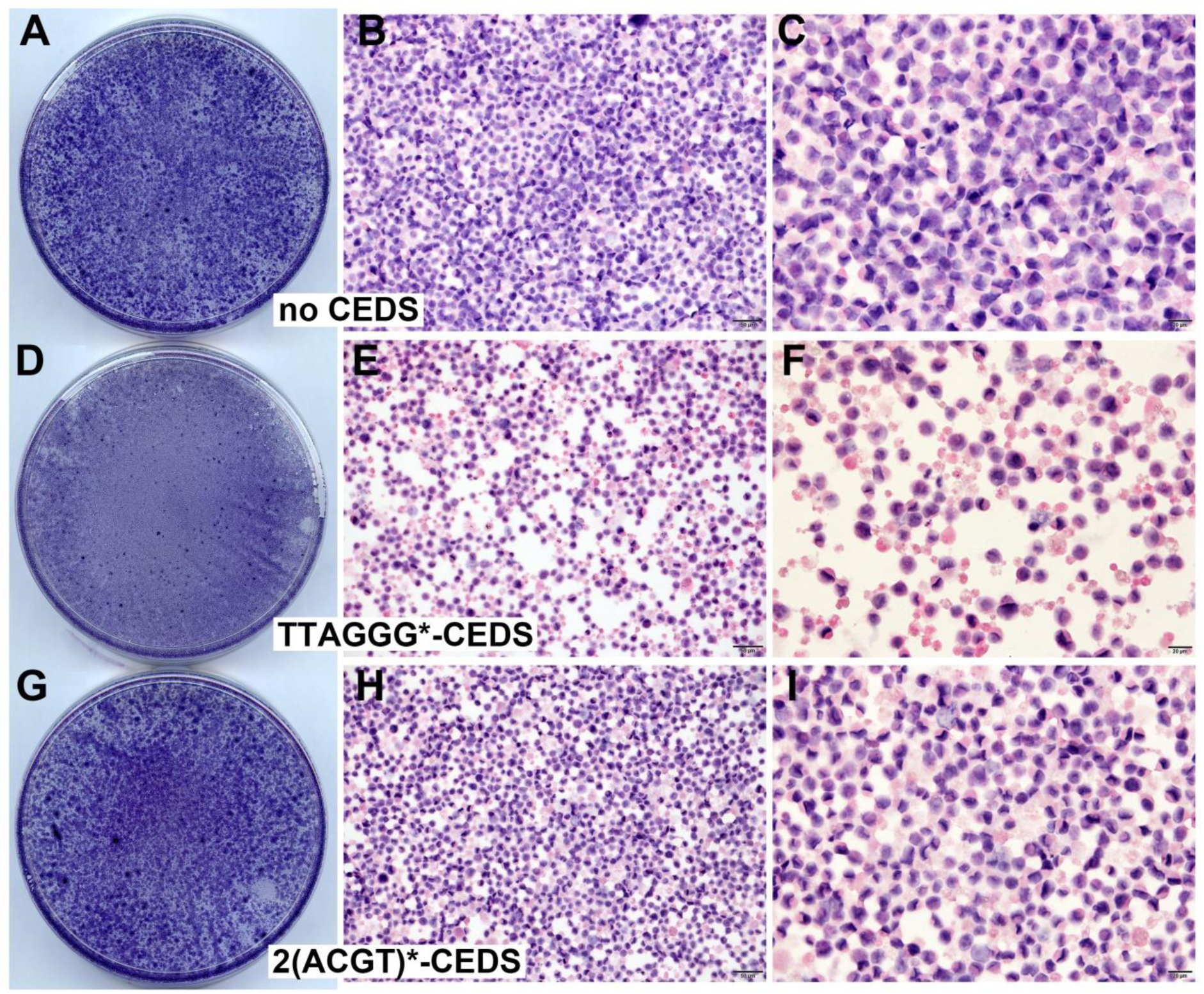
Cytological observation of KB cells. H&E stain. Visual and microscopic view of cells treated with TTAGGG*-CEDS (D-F), 2(ACGT)*-CEDS (G-I), or no CEDS (A-C). F: Noted severe apoptosis with increased empty space.

### Cytological observation of KB cells treated with CEDS using a telomere repeat sequence or a canonical E-box sequence

KB cells were cultured on plastic dishes as described above. Approximately 50% confluent KB cells grown on culture dish surfaces were treated with dodecagonal CEDS using a target sequence at 20-25 Gauss for 20 min twice a day in the culture incubator. Following three days of experiments, the cells were fixed with 10% buffered formalin and stained with toluidine blue, a basic thiazine metachromatic dye with high affinity for acidic tissue components, likely DNA and RNA. Subsequently, visual and microscopic observations were performed.

A comparison of CEDS effect between telomere repeat sequence*-CEDS (TTAGGG*-CEDS) and mutant telomere repeat sequence*-CEDS (TTTGGG*-CEDS) revealed that KB cells were significantly decreased in cell density and metachromasia in toluidine blue by TTAGGG*-CEDS compared to untreated control, while only a slight decrease by TTTGGG*-CEDS (Fig. 2 A-C). The results showed that TTAGGG*-CEDS strongly induced cell death, while TTTGGG*-CEDS induced cell death to a lesser extent than TTAGGG*-CEDS due to a base pair inversion from A-T to T-A. On the other hand, CACGTG*-CEDS induced almost no change in cell density, cell number, metachromasia in toluidine blue staining both visually and microscopically compared to 2(ACGT)*-CEDS and untreated control (Fig. 2 D-F).

**Figure 2.**
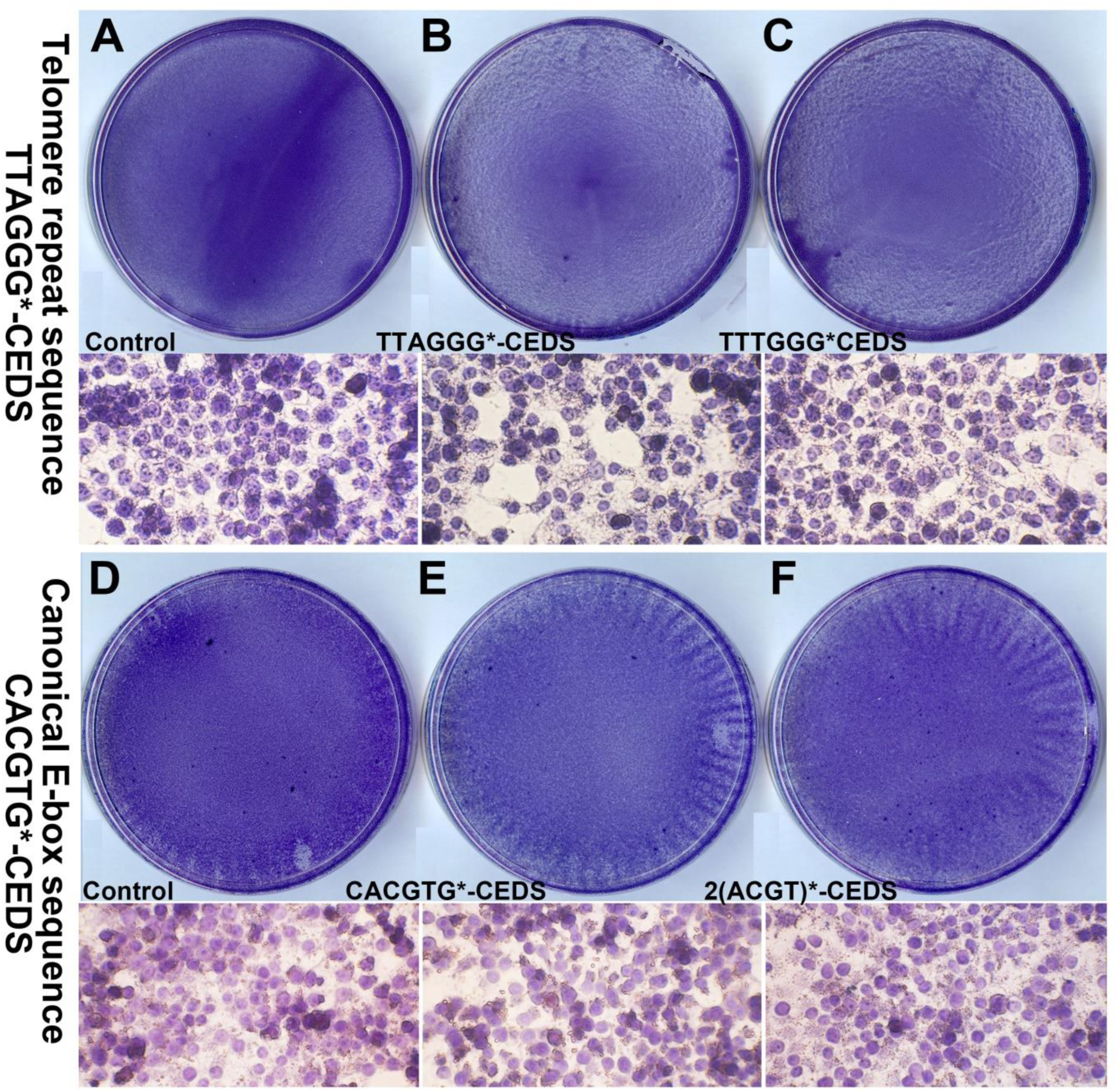
Cytological observation of KB cells treated with CEDS at 20-25 Gauss, twice daily for 20 min, during 3 days of cell culture with toluidine blue stain. For the comparison of CEDS-induced cytological results, TTAGGG*-CEDS and TTTGGG*-CEDS were assessed compared to no CEDS (A-C), and CACGTG*CEDS was compared with 2(ACGT)*-CEDS and no CEDS (D-F).

### IP-HPLC analysis for CEDS-induced changes of protein signaling pathways in RAW 264.7 cells

IP-HPLC has been developed to analyze the protein expression levels in comparison with the controls (*5, 6*). The immunoprecipitated proteins were subjected to the analysis with a HPLC (Agilent, USA) using a reverse phase column packed with non-adherent silica beads. The control and experimental samples were analyzed in sequence to permit a comparison of their HPLC peaks. IP-HPLC is available to use only small amount of total protein, up to 20-50 μg, thereby it can detect wide range of protein signaling simultaneously and repeatedly even with the limited amount of protein sample. However, IP-HPLC can provide the protein expression level in a simple, fast, accurate, cheap, multiple, and automatic way compared to other ordinary protein detection methods. In addition, the IP-HPLC data are available for statistical analysis.

The RAW 264.7 cells, derived from monocytes/macrophage-like cells of Balb/c mice, were utilized by limiting their culture passage number to less than 20 (*7*). The cells were cultured in Dulbecco’s modified Eagle’s medium supplemented with 10% fetal bovine serum, 100 units/mL carbenicillin, 100 μg/mL streptomycin, and 250 ng/mL amphotericin B (WelGene, Korea), in 5% CO_2_ incubator at 37.5°C. The cell culture product containing 10^8^-10^9^ cells was placed in the incubator and treated with dodecagonal CEDS using a target sequence at 20-25 Gauss, 5% CO2, and 37℃ for 20 minutes, left in the incubator for another 30 minutes, and harvested by centrifugation at 150g for 10 min for the following IP-HPLC assay.

The RAW 264.7 cells were lysed using protein lysis buffer (iNtRON Biotechnology, Korea), and analyzed by IP-HPLC as follow. Protein A/G agarose columns were separately pre-incubated with each 1 μg of 350 antibodies (Table S2). The supernatant of the antibody-incubated column was removed, and followed by IP-HPLC. Briefly, each protein sample, 50-100 μg, was mixed with 5 mL of binding buffer (150mM NaCl, 10mM Tris pH 7.4, 1mM EDTA, 1mM EGTA, 0.2mM sodium vanadate, 0.2mM PMSF and 0.5% NP-40) and incubated in the antibody-bound protein A/G agarose bead column on a rotating stirrer at room temperature for 1 h. After multiple washing of the columns with Tris-NaCl buffer, pH 7.5, in a graded NaCl concentration (0.15–0.3M), the target proteins were eluted with 300μL of IgG elution buffer (Pierce, USA). The immunoprecipitated proteins were analyzed using a precision HPLC unit (1100 series, Agilent, Santa Clara, CA, USA) equipped with a reverse-phase column and a micro-analytical UV detector system (SG Highteco, Hanam, Korea). Column elution was performed using 0.15M NaCl/20% acetonitrile solution at 0.5 mL/min for 15 min, 40°C, and the proteins were detected using a UV spectrometer at 280 nm. The control and experimental samples were run sequentially to allow comparisons.

For IP-HPLC, the whole protein peak areas (mAUs*) were obtained and calculated mathematically using an analytical algorithm by subtracting the negative control antibody peak areas, and protein expression levels were compared and normalized using the square roots of protein peak areas. IP-HPLC is available to use only small amount of total protein, up to 20-50 μg, thereby it can detect a wide range of protein signaling simultaneously and repeatedly even with the limited amount of protein sample. Since IP-HPLC gives the relative ratio (%) of protein expression compared to untreated control, the protein expression data could be analyzed statistically and widely compared with other protein expressions. The ratios were divided into five categories; severe underexpression (below 80%), marked underexpression (80%-below 95%), minimal change (95%-below 105%), marked overexpression (105%-120%), and severe overexpression (above 120%).

Although the immunoprecipitation is unable to define the size-dependent expression of target protein compared to western blot, it collects every protein containing a specific epitope against antibody in the protein samples from cells, blood, urine, saliva, inflammatory exudate, etc. (*5, 6, 8–17*). Therefore, IP-HPLC can detect whole precursor and modified target proteins similar to enzyme-linked immunosorbent assay (ELISA), but the antigen-antibody binding strength can be adjusted by different salt buffers depending on the condition of the protein samples.

### Telomere repeat sequence*-CEDS induced potent anticancer signaling in RAW 264.7 cells

IP-HPLC results showed that TTAGGG*-CEDS upregulated 34 proteins (31.6%), downregulated 67 proteins (62.6%), and exhibited minimum effect on six proteins (5.6%) out of 107 telomere-associated proteins observed in this study. In addition, The TTAGGG*-CEDS affected the overall protein signaling pathways in RAW 264.7 cells by downregulating telomere-associated proteins and reactive proteins as well (Fig. 3A, Table 1), which were summarized as follows according to the expression of telomere-associated proteins.

**Figure 3.**
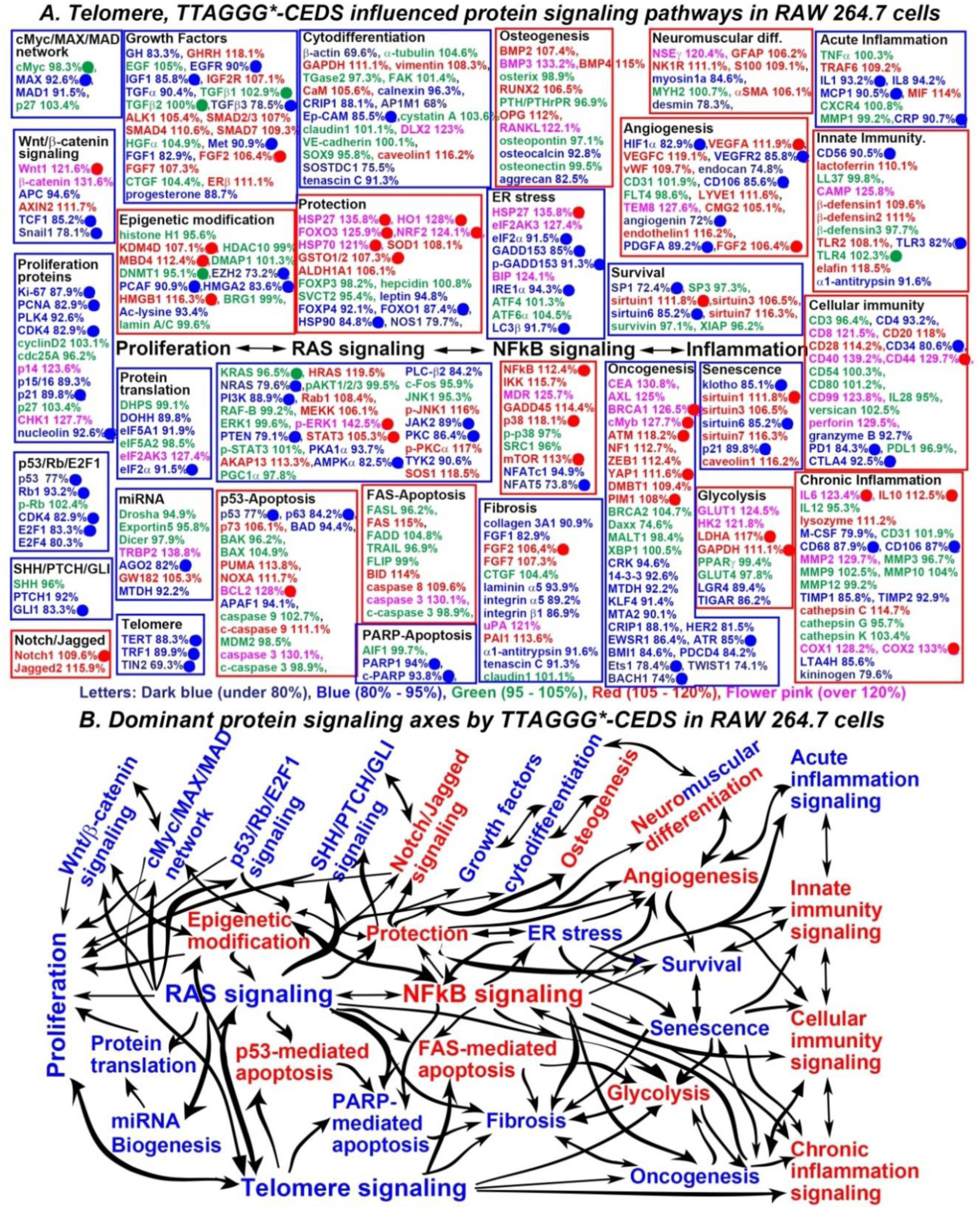
TTAGGG*-CEDS influenced the protein signaling pathways (A) and axes (B) in RAW 264.7 cells. In IP-HPLC, proteins downregulated (blue), upregulated (red), and minimally changed (green) compared to untreated controls. Dominantly suppressed (blue square) and activated (red square) signaling. Telomere-associated proteins (Harmonizome 3.0) were downregulated (●) or upregulated (●) by TTAGGG*-CEDS.

**Table 1.** Protein expressions in RAW 264.7 cells treated with telomere sequence (TTAGGG)*-CEDS.

| Signaling pathways | Downregulated Proteins | Upregulated Proteins |
| --- | --- | --- |
| Proliferation | Ki-67, PCNA, PLK4, CDK4, p15/16, p21, nucleolin | p14, CHK1 |
| cMyc/MAX/MAD network | MAX, MAD1 |  |
| Wnt/β-catenin signaling | APC, TCF1, Snail | Wnt1, β-catenin, AXIN2 |
| p53/Rb/E2F1 signaling | p53, Rb1, CDK4, E2F1, E2F4 |  |
| SHH/PTCH/GLI signaling | PTCH1, GLI1 |  |
| Notch/Jagged signaling |  | Notch1, Jagged2 |
| Epigenetic modification | EZH2, PCAF, HMG2, Ac-lysine | KDM4D, MBD4, HMGB1 |
| Protein translation | DOHH, eIF5A1, eIF2α | eIF2AK3 |
| miRNA biogenesis | AGO2, MTDH | TRBP2, GW182 |
| Growth factors | GH, EGFR, IGF1, TGFα, TGFβ3, Met, FGF1, progesterone | GHRH, IGF2R, ALK1, SMAD2/3, SMAD4, SMAD7, HGFα, FGF2, FGF7, ERβ |
| Cytodifferentiation | β-actin, calnexin, CRIP1, AP1M1, Ep-CAM, SOSTDC1, tenascin C | GAPDH, vimentin, CaM, DLX2, caveolin1 |
| Osteogenesis | osteocalcin, aggrecan | BMP2, BMP3, BMP4, RUNX2, OPG, RANKL |
| Neuromuscular differentiation | myosin-1a, desmin | NSEγ, GFAP, NK1R, S100, αSMA |
| RAS signaling | NRAS, PI3K, PTEN, PKA1α, AMPKα, PLCβ2, JAK2, PKC, TYK2 | HRAS, Rab1, MEKK, p-ERK1, STAT3, AKAP13, p-JNK1, p-PKC1α, SOS1 |
| NFκB signaling | SRC1, NFATc1, NFAT5 | NFκB, IKK, MDR, GADD45, p38, mTOR |
| p53-mediated apoptosis | p53, p63, BAD, APAF1 | p73, BAK, PUMA, NOXA, BCL2, c-caspase9, caspase3 |
| FAS-mediated apoptosis | FASL | FAS, FADD, BID, caspase8, caspase3 |
| PARP-mediated apoptosis | PARP1, c-PARP |  |
| Protection | leptin, FOXP4, FOXO1, HSP90, NOS1 | HSP27, HO1, FOXO3, NRF2, HSP70, SOD1, GSTO1/2, ALDH1A1 |
| ER stress | eIF2α, GADD153, p-GADD153, IRE1α, LC3β | HSP27, eIF2AK3, BIP |
| Angiogenesis | HIF1α, VEGFR2, endocan, CD106, angiogenin, PDGFA | VEGFA, VEGFC, vWF, LYVE1, TEM8, CMG2, endothelin1, FGF2 |
| Survival | SP1, sirtuin6 | sirtuin1, sirtuin3, sirtuin7 |
| Acute inflammation | IL1, IL8, MCP1, CRP | TRAF6, MIF |
| Innate immunity | CD56, TLR3, α1-antitrypsin | Lactoferrin, CAMP, β-defensin1, β-defensin2, TLR2, elafin |
| Cellular immunity | CD4, CD34, granzyme B, PD1, CTLA4 | CD8, CD20, CD28, CD40, CD44, CD99, perforin |
| Chronic inflammation | M-CSF, CD68, CD106, TIMP1, TIMP2, LTA4H, kininogen | IL6, IL10, lysozyme, MMP2, cathepsin C, COX1, COX2 |
| Glycolysis | LGR4, PPARγ, TIGAR, | GLUT1, HK2, LDHA, GAPDH |
| Fibrosis | collagen 3A1, FGF1, laminin α5, integrin α5, integrin β1, α1-antitrypsin, tenascin C | FGF2, FGF7, uPA, PAI1 |
| Oncogenesis | CRK, 14-3-3, MTDH, KLF4, MTA2, CRIP1, HER2, EWSR1, ATR, BMI1, PDCD4, Ets1, TWIST1, BACH1 | CEA, AXL, BRCA1, cMyb, ATM, NF1, ZEB1, YAP1, DMBT1, PIM1 |
| Senescence | klotho, sirtuin6, p21 | sirtuin1, sirtuin3, sirtuin7, caveolin1 |
| Telomere signaling | TERT, TRF1, TIN2 |  |

***Proliferating signaling*** was suppressed by downregulating Ki-67 (87.9%), PCNA (82.9%), CDK4 (82.9%), p21 (89.8%), nucleolin (92.8%). ***cMyc/MAX/MAD network*** was suppressed by downregulating MAX (92.8%). ***Wnt/β-catenin signaling*** was suppressed by downregulating TCF1 (85.2%), Snail1 (78.1%).

***P53/Rb/E2F1 signaling*** was suppressed by downregulating p53 (77%), Rb1 (93.2%), CDK4 (82.9%), E2F1 (83.3%). ***Protein translation signaling*** was suppressed by downregulating DOHH (89.8%), eIF2α (91.5%). ***MiRNA biogenesis signaling*** was suppressed by downregulating AGO2 (82%).

***Growth factor signaling*** was partially suppressed by downregulating EGFR (90%), IGF1 (85.8%), TGFβ3 (78.5%), Met (90.9%). ***Cytodifferentiation signaling*** was partially suppressed by downregulating Ep-CAM (85.5%). ***RAS signaling*** was suppressed by downregulating, NRAS (79.6%), PI3K (88.9%), PTEN (79.1%), AMPKα (82.5%), JAK2 (89%), PKC (86.4%).

***ER stress*** was suppressed by downregulating eIF2α (91.5%), GADD153 (85%), p-GADD153 (91.3%), IRE1α (94.3%), LC3β (91.7%). ***Survival signaling*** was suppressed by downregulating SP1 (72.4%), sirtuin6 (85.2%). ***Acute inflammation signaling*** was suppressed by downregulating IL1 (93.2%), MCP1 (90.5%), CRP (90.7%). ***PARP-mediated apoptosis signaling*** was suppressed by downregulating, PARP1 (94%), c-PARP (93.8%).

***Fibrosis signaling*** was suppressed by downregulating collagen 3A1 (90.9%), FGF1 (82.9%), laminin α5 (93.9%), integrin α5 (89.2%), integrin β1 (86.9%), α1-antitrypsin (91.6%), tenascin C (91.3%). ***Senescence signaling*** was suppressed by downregulating klotho (85.1%), sirtuin6 (85.2%), p21 (89.8%). ***Telomere signaling*** was suppressed by downregulating TERT (88.3%), TRF1 (89.9%), TIN2 (69.3%).

Conversely, ***Notch/Jagged signaling*** was activated by upregulating Notch1 (109.6%). ***Osteogenesis signaling*** appears to be activated by upregulating BMP2 (107.4%), BMP3 (133.2%), BMP4 (115%), RUNX2 (106.5%), OPG (112%), RANKL (122.1%). ***Neuromuscular differentiation signaling*** was activated by upregulating NSEγ (120.4%), GFAP (106.2%), NK1R (111.1%), S100 (109.1%), αSMA (106.1%).

***NFkB signaling*** was activated by upregulating NFkB (112.4%), p38 (118.1%), mTOR (113%). ***Protection signaling*** was activated by upregulating HSP27 (135.8%), HO1 (128%), FOXO3 (125.9%), NRF2 (124.1%), HSP70 (121%), GSTO1/2 (107.3%). ***Angiogenesis signaling*** was activated by upregulating VEGFA (111.9%), FGF2 (106.4%). ***Innate immunity signaling*** appears to be activated by upregulating lactoferrin (110.1%), CAMP (125.8%), β-defensin1 (109.6%), β-defensin2 (111%), TLR2 (108.1%), elafin (118.5%). ***Cellular immunity signaling*** was activated by upregulating CD44 (129.7%).

***Chronic inflammation signaling*** was activated by upregulating IL6 (123.4%), IL10 (112.5%), COX2 (133%). ***P53-mediated apoptosis signaling*** appears to be activated by upregulating p73 (106.1%), PUMA (113.8%), NOXA (111.7%), c-caspase9 (111.1%), caspase3 (130.3%). ***FAS-mediated apoptosis signaling*** was activated by upregulating FAS (115%), BID (114%), caspase8 (109.6%), caspase3 (130.1%). ***Glycolysis signaling*** was activated by upregulating LDHA (117%), GAPDH (111.1%).

In addition, TTAGGG*-CEDS variably influenced ***epigenetic modification signaling*** by upregulating proteins involved in DNA methylation/acetylation such as KDM4D (107.1%), MBD4 (112.4%), HMGB1 (116.3%), therefore, it induced a trend towards an increase in the methylation of histones and DNAs, transcriptional repression. TTAGGG*-CEDS broadly affected ***oncogenesis signaling*** by upregulating such as BRCA1 (126.5%), cMyb (127.7%), ATM (118.2%), YAP1 (111.6%), PIM1 (108%), significantly impacting the entire protein signaling pathways in RAW 264.7 cells. In addition to the expressions of telomere-associated proteins, the expressions of telomere-unassociated proteins in the cells treated with TTAGGG*-CEDS showed similar trends to the above protein signaling (Fig. 3, Table 1).

Resultantly, TTAGGG*-CEDS primary inhibited *RAS signaling*, subsequently inactivating *protection-ER stress-survival signaling axis, p53– and FAS-mediated apoptosis signaling axis,* and *telomere-oncogenesis-PARP-associated apoptosis-fibrosis axis*, while activated *NF-kB signaling*, subsequently stimulating epigenetic modification-protection-angiogenesis signaling axis, p53– and FAS-mediated apoptosis-glycolysis-chronic inflammation signaling axis *(Fig. 3B)*.

The results indicate that TTAGGG*-CEDS induced potent anti-oncogenic effect with extensive apoptosis on RAW 264.7 cells by inhibiting cell proliferation, ER stress, survival, cellular senescence, telomere instability, and oncogenesis, and concurrently enhancing ROS protection, angiogenesis, apoptosis, glycolysis, and chronic inflammation. It is noteworthy that TTAGGG*-CEDS also has the potential to enhance innate and cellular immunity.

### Canonical E-box sequence*-CEDS induced significant oncogenic signaling in RAW 264.7 cells

IP-HPLC results showed that CACGTG*-CEDS upregulated 57proteins (35.8%), downregulated 75 proteins (47.2%), and exhibited minimum effect on 27 proteins (17%) out of 159 cMyc target proteins. In addition, CACGTG*-CEDS concomitantly upregulated 77 proteins (40.3%) and downregulated 58 proteins (30.4%) out of 191 cMyc non-target proteins indirectly. The CACGTG*-CEDS affected the overall protein signaling pathways in RAW 264.7 cells by downregulating cMyc target proteins and reactive proteins as well (Fig. 4A, Table 2), which were summarized as follows according to the expression of cMyc target proteins.

**Figure 4.**
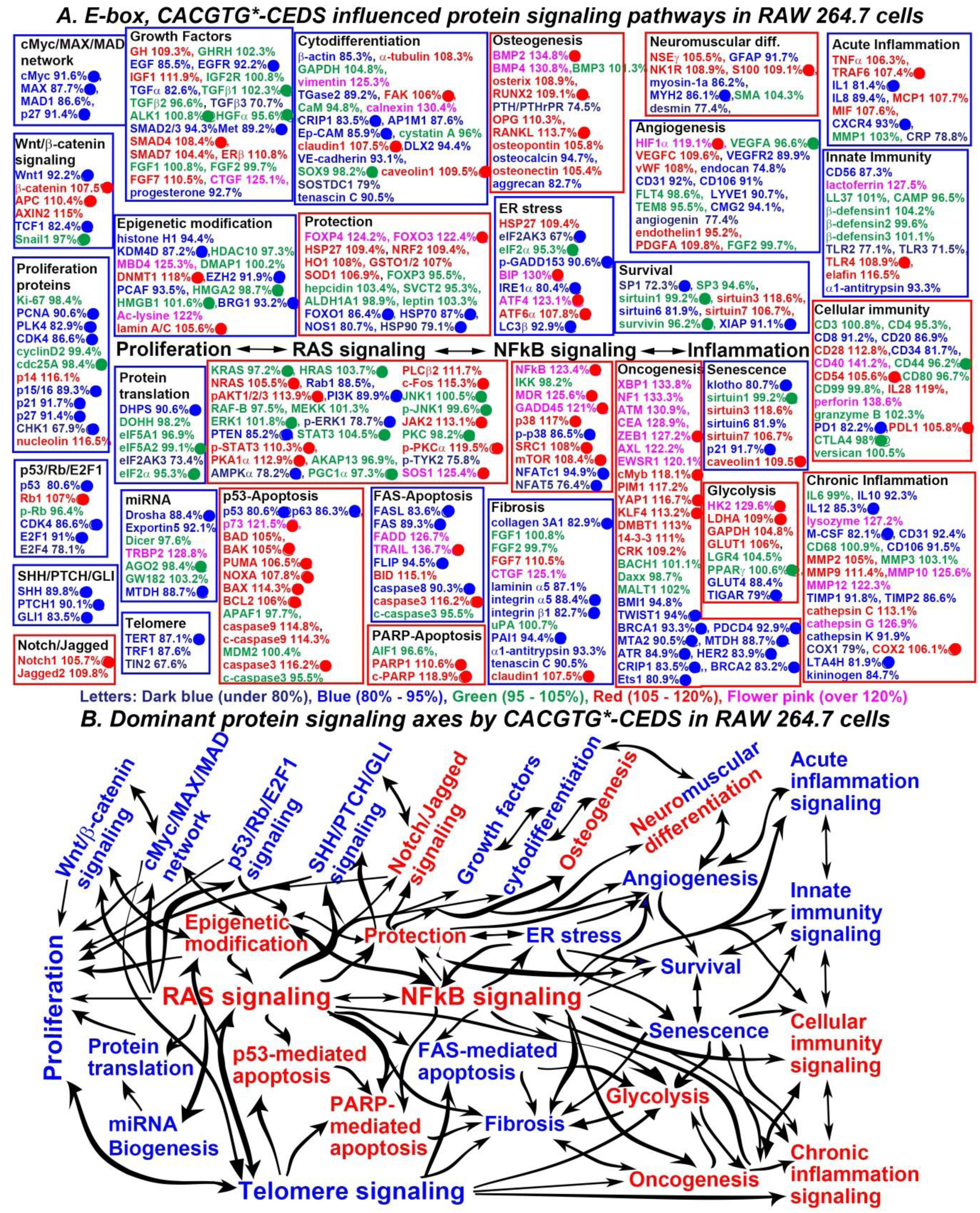
CACGTG*-CEDS influenced the protein signaling pathways (A) and axes (B) in RAW 264.7 cells. In IP-HPLC, proteins downregulated (**blue**), upregulated (**red**), and minimally changed (**green**) compared to untreated controls. Dominantly suppressed (**blue square**) and activated (**red square**) signaling. cMyc target proteins (Harmonizome 3.0) were downregulated (●) or upregulated (●) by CACGTG*-CEDS.

**Table 2.**
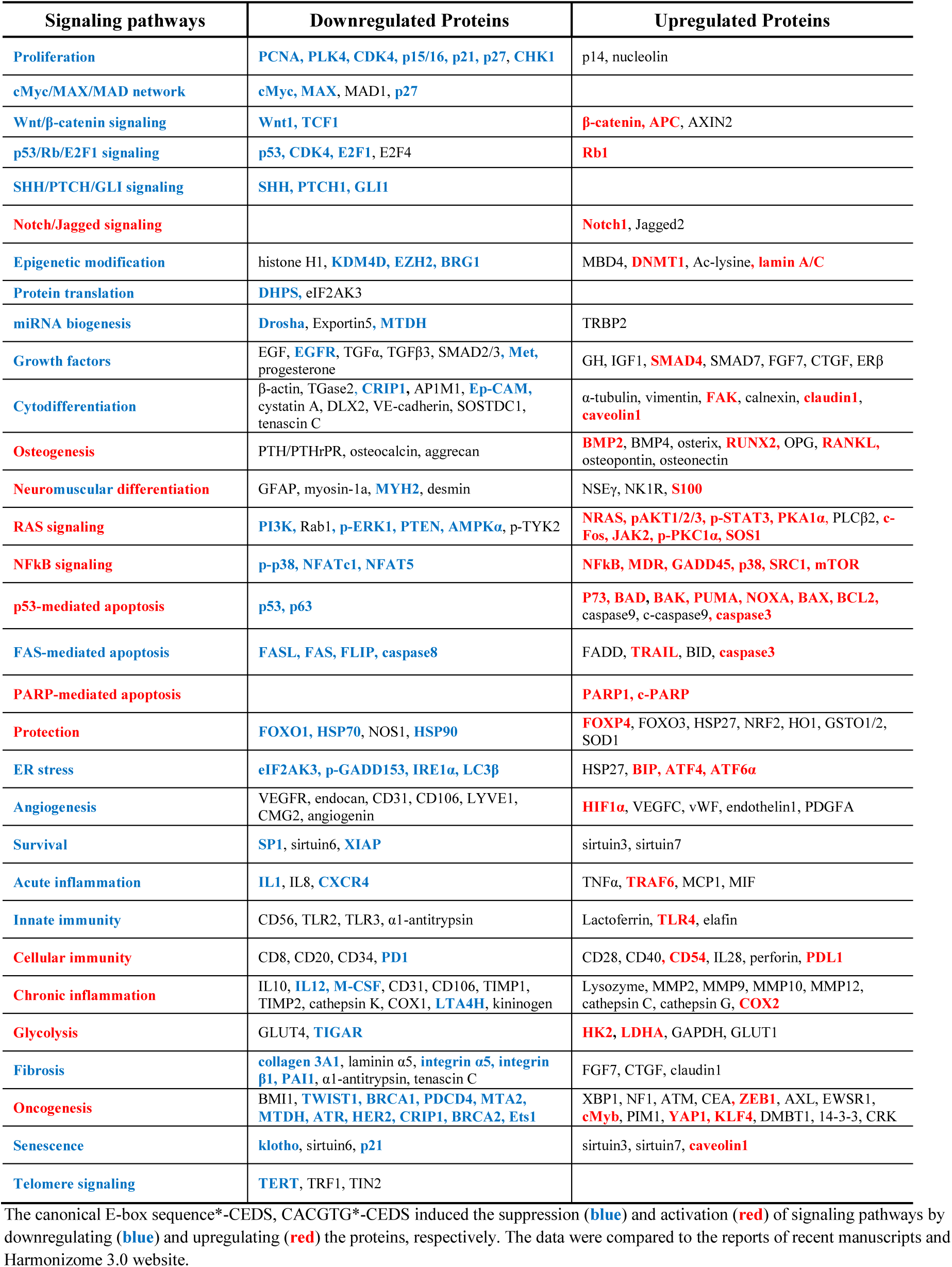
Protein expressions in RAW 264.7 cells treated with canonical E-box sequence (CACGTG)*-CEDS.

***Proliferating signaling*** was partially inactivated by downregulating PCNA (90.6%), PLK4 (82.9%), CDK4 (86.6%). ***cMyc/MAX/MAD network*** was suppressed by downregulating cMyc (91.6%), MAX (87.7%), p27 (91.4%). ***Wnt/β-catenin signaling*** appears to be inactivated by downregulating Wnt1 (92.2%), TCF1 (82.4%).

***P53/Rb/E2F1 signaling*** was suppressed by downregulating p53 (80.6%), CDK4 (86.6%), E2F1 (91%). ***SHH/PTCH/GLI signaling*** was suppressed by downregulating SHH (89.8%), PTCH1 (90.1%), GLI (83.5%). ***Protein translation signaling*** appears to be inactivated by downregulating DHPS (90.6%). ***miRNA biogenesis signaling*** was inactivated by downregulating Drosha (88.4%), MTDH (88.7%).

***Growth factor signaling*** was partially inactivated by downregulating, EGFR (92.2%), Met (89.2%). ***Cytodifferentiation signaling*** was partially suppressed by downregulating CRIP1 (83.5%), Ep-CAM (85.9%). ***ER stress*** was inactivated by downregulating eIF2AK3 (67%), p-GADD153 (90.6%), IRE1α (90.4%), LC3β (92.9%). ***Angiogenesis signaling*** appears to be inactivated by downregulating VEGFR2 (89.9%), endocan (74.8%), CD31 (92%), CD106 (91%), LYVE1 (90.7%), CMG2 (94.1%), angiogenin (77.4%).

***Survival signaling*** was suppressed by downregulating SP1 (72.3%), XIAP (91.1%). ***Acute inflammation signaling*** appears to be inactivated by downregulating IL1 (81.4%), CXCR4 (93%). ***Innate immunity signaling*** appears to be inactivated by downregulating CD56 (87.3%), TLR2 (77.1%), TLR3 (71.5%), α1-antitrypsin (93.3%). ***Chronic inflammation signaling*** was inactivated by downregulating IL12 (85.3%), M-CSF (82.1%), LTA4H (81.9%).

***FAS-mediated apoptosis signaling*** was suppressed by downregulating FASL (83.6%), FAS (89.3%), FLIP (94.5%), caspase8 (90.3%). ***Fibrosis signaling*** was suppressed by downregulating collagen 3A1 (82.9%), integrin α5 (88.4%), integrin β1 (82.7%), PAI1 (94.4%). ***Senescence signaling*** appears to be suppressed by downregulating klotho (80.7%), p21 (91.7%), and coincidentally downregulating sirtuin6 (81.9%). ***Telomere signaling*** was suppressed by downregulating TERT (87.1%), TRF1 (87.6%), TIN2 (67.6%).

Conversely, ***Notch/Jagged signaling*** was activated by upregulating Notch1 (105.7%). ***Osteogenesis signaling*** was activated by upregulating BMP2 (134.8%), RUNX2 (109.1%), RANKL (113.7%). ***Neuronal differentiation signaling*** was activated by upregulating S100 (109.1%), while ***muscular differentiation signaling*** was inactivated by downregulating MYH2 (86.1%).

***RAS signaling*** appears to be activated by upregulating NRAS (105.5%), pAKT1/2/3 (113.9%), p-STAT3 (110.3%), PKA1α (112.9%), c-Fos (115.3%), JAK2 (113.1%), p-PKCα (119.5%), SOS1 (125.4%). ***NFkB signaling*** was activated by upregulating NFkB (123.4%), MDR (125.6%), GADD45 (121%), p38 (117%), SRC1 (108%), mTOR (108.4%). ***Protection signaling*** appears to be activated by upregulating FOXO3 (122.4%).

***Cellular immunity signaling*** appears to be activated by upregulating CD54 (105.6%) and downregulating PD1 (82.2%). ***p53-mediated apoptosis signaling*** was activated by upregulating p73 (121.5%), BAK (105%), PUMA (106.5%), NOXA (107.8%), BAX (114.3%), BCL2 (106%), caspase3 (116.2%). ***PARP-mediated apoptosis signaling*** was activated by upregulating PARP1 (110.6%), c-PARP (118.9%). ***Glycolysis signaling*** appears to be activated by upregulating HK2 (129.6%), LDHA (109%).

In addition, CACGTG*-CEDS, markedly influenced ***epigenetic modification signaling*** by downregulating KDM4D (87.2%), EZH2 (91.9%), BRG1 (93.2%), significantly suppressed ***oncogenesis signaling*** by downregulating TWIST (94%), BRCA1 (93.3%), PDCD4 (92.9%), MTA2 (90.5%), MTDH (88.7%), ATR (84.9%), HER2 (83.9%), CRIP1 (83.5%), BRCA2 (83.2%), Ets1 (80.9%). Besides the expressions of cMyc target proteins, the expressions of cMyc non-target proteins in the cells treated with CACGTG*-CEDS were summarized in Table S3.

Consequently, CACGTG*-CEDS strongly impacted the entire protein signaling pathways in RAW 264.7 cells. CACGTG*-CEDS enhanced ***RAS-NFkB signaling axis***, subsequently activating ***epigenetic modification-protection-Notch/Jagged-osteogenesis signaling axis, p-53 and PARP-mediated apoptosis signaling axis, glycolysis-oncogenesis-chronic inflammation,*** and ***cellular immunity signaling axis***, while it inactivated ***proliferation-related signaling axis, ER stress-angiogenesis-survival-senescence-acute inflammation-innate immunity signaling axis,*** and ***FAS-mediated apoptosis-fibrosis-telomere signaling axis*** (Fig. 4B).

The results indicate that CACGTG*-CEDS increased the oxidative stress, p53– and PARP-mediated apoptosis, glycolysis, oncogenesis, and chronic inflammation in RAW 264.7 cells, but decreased the proliferation, ER stress, angiogenesis, survival, senescence, acute inflammation, FAS-mediated apoptosis, and telomere instability, thereby, providing oncogenic environment with reduced potential of the cytodifferentiation, regeneration, wound healing, and innate and cellular immunity in the cells.

### Nonspecific poly-A 12A*-CEDS induced imbalanced protein signaling in RAW 264.7 cells

RAW 264.7 cells were also treated with CEDS using nonspecific poly-A sequence (12A*-CEDS) as a positive control, and analyzed by IP-HPLC to compared with the results of TTAGGG*-CEDS and CACGTG*-CEDS. In constrast, 12A*-CEDS resulted in imbalanced protein signaling in RAW 264.7 cells, activating epigenetic methylation, oncogenesis, and telomere instability, leading to chronic inflammation and apoptosis, in the absence of cellular growth and differentiation, wound healing, and ROS protection (Fig. 5, Table 4). The variable protein expressions after 12A*-CEDS were detected by IP-HPLC using 350 antisera, and their imbalaced influences in different signaling pathways were summarized and described in Supplement Text.

**Figure 5.**
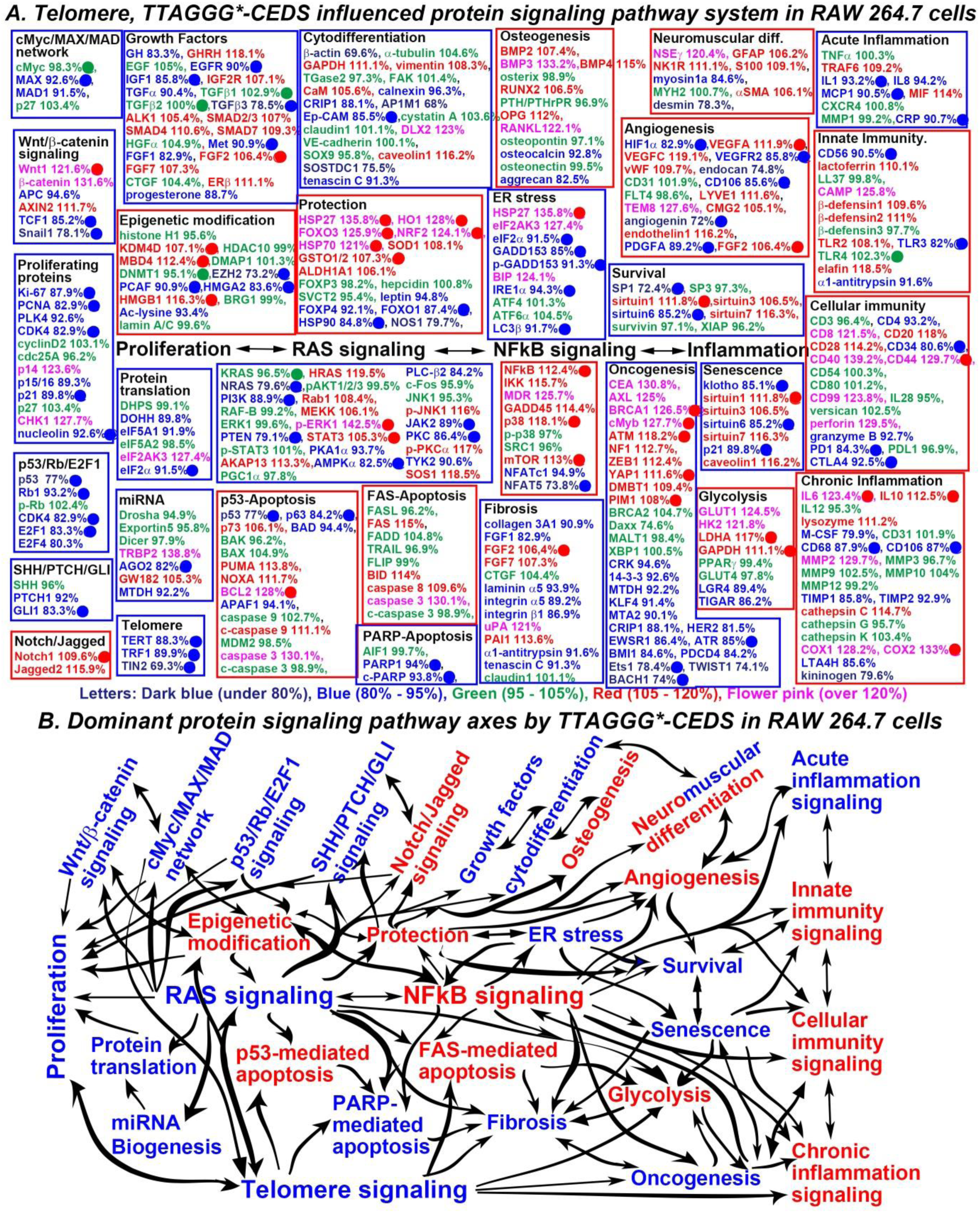
The telomere sequence (TTAGGG)*-CEDS influenced the protein signaling pathway system (A) and axes (B) in RAW 264.7 cells. The IP-HPLC results revealed proteins that were downregulated (blue), upregulated (red), and minimally changed (green) compared to the untreated controls. Dominantly suppressed (blue square) and activated (red square) signaling pathways. Telomere-associated proteins (Harmonizome 3.0), downregulated (●) or upregulated (●) by the TTAGGG*-CEDS.

**Table 3.** The targeting efficiency of CEDS using a different DNA motif sequence in RAW 264.7 cells.

| **Protein binding site**<br>**sequence\*-CEDS** | **Up-regulation**<br>**(over 105%)** | **Down-**<br>**regulation**<br>**(under 95%)** | **Minimal**<br>**change**<br>**($\pm 5\%$)** | **Total**<br>**target**<br>**proteins** |
| --- | --- | --- | --- | --- |
| **TTAGGG\*-CEDS** | 34 | 67 | 6 | 107 |
| **(telomere repeat sequence)** | 31.6% | 62.6% | 5.6% | 100% |
| **CACGTG\*-CEDS** | 57 | 75 | 27 | 159 |
| **(canonical E-box sequence)** | 35.8% | 47.2% | 17% | 100% |
| **Total** | 91 | 142 | 33 | 266 |
|  | 34.2% | 53.4% | 12.4% | 100% |
| **12A\*-CEDS** | 130 | 122 | 98 | 350 |
| **(nonspecific sequence)** | 37.1% | 34.9% | 28% | 100% |

**Table 4.**
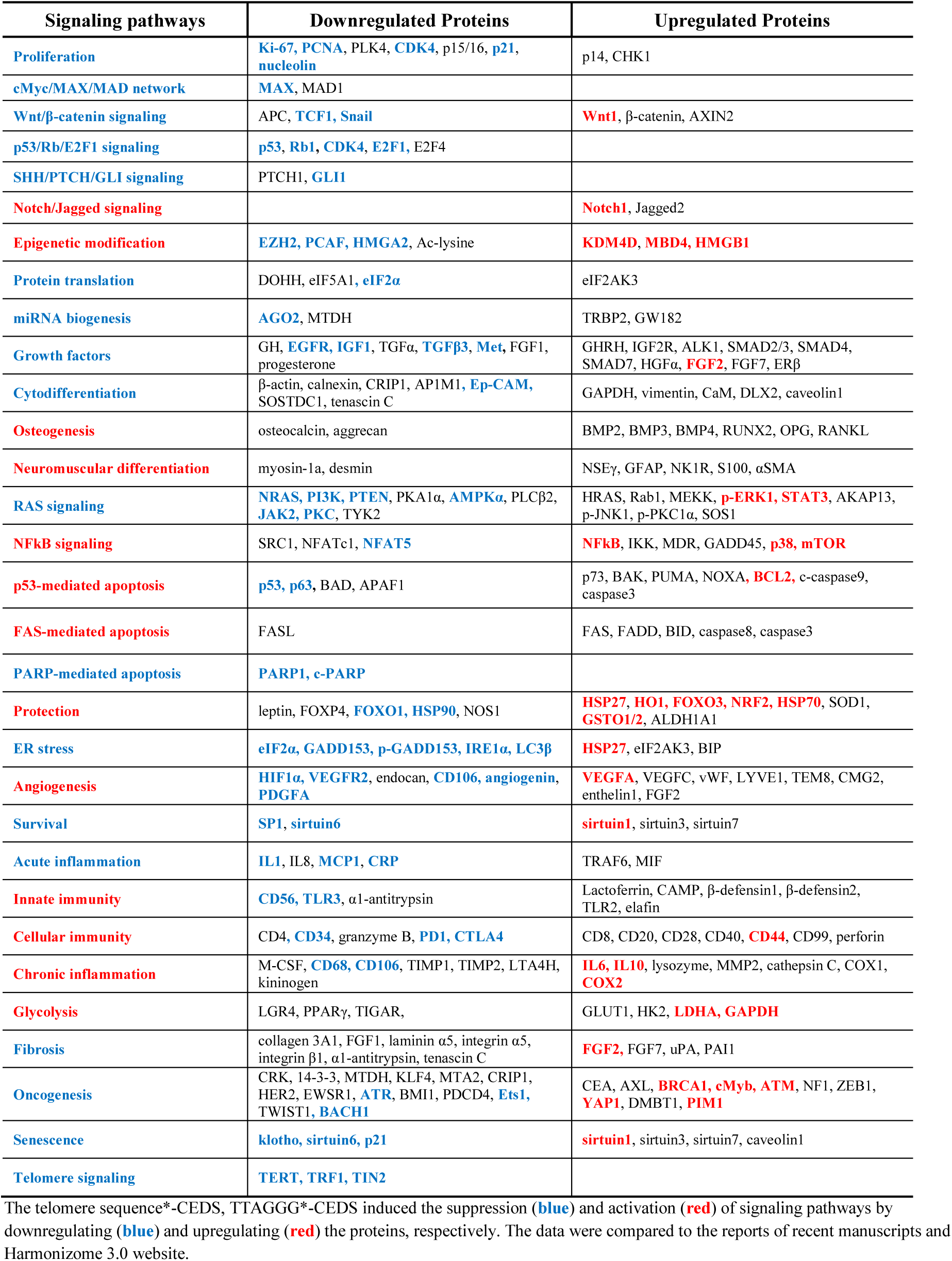
Protein expressions in RAW 264.7 cells treated with telomere sequence (TTAGGG)*-CEDS.

Therefore, it is postulated that 12A*-CEDS induced irregular upregulation and downregulation of multiple signaling pathways due to imbalanced stimulation of gene transcription by targeting poly-A sequences widely distributed in genome, resulted in the aging and retrogressive changes with a high potential of oncogenesis.

## Discussion

This study demonstrated the CEDS targeting a six bps telomere repeat sequence and a six bps palindromic canonical E-box sequence in RAW 264.7 cells. Both sequences are favorable as a CEDS target due to their abundant distribution throughout the genome and their important roles in gene transcription and regulation (*4, 18*). As a result, TTAGGG*-CEDS inactivated telomere signaling, RAS-proliferation-growth-cytodifferentiation signaling axis, while enhancing NFkB-ROS-inflammation signaling axis, p53– and FAS-mediated apoptosis signaling with concomitant inactivation of ER stress-survival-senescence signaling axis, resulting in extensive cell death as seen in the cytological observation.

However, it was found that TTAGGG*-CEDS upregulated 34 proteins (31.6%), downregulated 67 proteins (62.6%), and had minimal effect on 6 proteins (5.6%) out of 107 telomere-associated proteins observed in this study. Since TTAGGG*-CEDS tended to downregulate rather than upregulate the telomere-associated proteins, it is postulated that TTAGGG*-CEDS stabilizes the telomere sequences, thereby, inactivates the telomere-associated proteins and subsequently attenuates the signaling relevant to telomere instability in cells.

In contrast, CACGTG*-CEDS activated RAS-NFkB signaling axis and subsequently enhanced ROS-p53 and PARP-associated apoptosis signaling, glycolysis-oncogenesis-chronic inflammation axis, while inactivated proliferation-growth axis, ER stress-angiogenesis-survival-senescence axis, resulting in the increased oncogenic potential and delayed regeneration and wound healing.

However, CACGTG*-CEDS upregulated 57 proteins (35.8%), downregulated 75 proteins (47.2%), and showed minimal effect on 27 proteins (17%) out of 159 cMyc target proteins, indicating relatively even upregulation (35.8%) and downregulation (47.2%) of cMyc target proteins compared to TTAGGG*-CEDS. Therefore, it is postulated that CACGTG*-CEDS affects palindromic six bps dsDNA, canonical dsCACGTG consisting of three Pyu dsDNA segments, by activating forward and reverse direction, leading to dual protein binding and sliding on the canonical E-box site. This may result in the selective upregulation and downregulation of cMyc target proteins depending on the context of molecular interaction, such as cMyc-MAX-MAD network.

Consequently, TTAGGG*-CEDS induced potent anticancer effect on RAW 264.7 cells by stabilizing the telomere sequence, which functions as a protective role for chromosome, while CACGTG*-CEDS induced significant oncogenic effect on the cells by activating the major oncogene, c-Myc. Therefore, it is suggested that CEDS can properly target DNA motifs such as telomere repeat sequence and canonical E-box sequence in the genomic DNA of RAW 264.7 cells, and upon further investigation, the TTAGGG*-CEDS and CACGTG*-CEDS will be useful candidate tools to regulate a specific protein signaling axis in cells.

Taken together, this series of study found that CEDS can target DNAs and RNAs distributed *in vitro* and *in vivo* environment in a sequence-specific manner. However, the CEDS-induced modulation of the target gene remains to be elucidated by further precise molecular genetic investigation.

## Acknowledgments

We would like to express our gratitude to the late Professor Je Geun Chi and the late Dr. Soo Il Chung, who contributed to this research in part.

